# The core transcriptome of mammalian placentas and the divergence of expression with placental shape

**DOI:** 10.1101/137554

**Authors:** Don L. Armstrong, Michael R. McGowen, Amy Weckle, Priyadarshini Pantham, Jason Caravas, Dalen Agnew, Kurt Benirschke, Sue Savage-Rumbaugh, Eviatar Nevo, Chong J. Kim, Günter P. Wagner, Roberto Romero, Derek E. Wildman

## Abstract

**Introduction:** The placenta is arguably the most anatomically variable organ in mammals even though its primary function is conserved.

**Method:** Using RNA-Seq, we measured the expression profiles of 55 term placentas of 14 species of mammals representing all major eutherian superordinal clades and marsupials, and compared the evolution of expression across clades.

**Results:** We identified a set of 115 core genes which is expressed (FPKM *≥* 10) in all eutherian placentas, including genes with immune-modulating properties (*ANXA2*, *ANXA1*, *S100A11*, *S100A10*, and *LGALS1*), cell-cell interactions (*LAMC1*, *LUM*, and *LGALS1*), invasion (*GRB2* and *RALB*) and syncytialization (*ANXA5* and *ANXA1*). We also identified multiple pre-eclampsia associated genes which are differentially expressed in *Homo sapiens* when compared to the other 13 species. Multiple genes are significantly associated with placenta morphology, including *EREG* and *WNT5A* which are both associated with placental shape.

**Discussion:** 115 genes are important for the core functions of the placenta in all eutherian species analyzed. The molecular functions and pathways enriched in the core placenta align with the evolutionarily conserved functionality of the placenta.

## 1. Introduction

Even though the placenta’s primary function is conserved among mammals, the placenta varies widely in terms of shape, degree of intimacy between fetal tissues and maternal tissues, and degree of interdigitation of maternal and fetal tissues in different species of mammals [1] (Supplemental Table S1). Mammals also differ drastically in the number and size of their offspring, as well as the length of gestation; these traits are often associated with specific aspects of placental morphology [2] (Supplemental Table S2). Although the morphology of the mammalian placenta has been well characterized in a number of species [1], the diversity of the molecular environment is only beginning to be understood from a genetic perspective [3].

These observations lead to two related questions: 1) what are the genes which are responsible for the **core functions of the placenta** and 2) which genes are associated with **changes in placenta morphology**?

Previous studies of some species (i.e., *Homo sapiens*, *Mus musculus*, *Bos taurus*, and *Loxodonta africana*) have provided some preliminary answers to these questions. They revealed a stunning divergence in the genetics of the placenta, especially in later stages of pregnancy [3–6]. For example, each superorder (Euarchontoglires, Laurasiatheria, Afrotheria, and Xenartha, Fig. 1) has evolved expansive placenta-specific gene families with expression patterns that can vary in space and time [7]. These include independent expansions of the prolactins in *Mus musculus*, *Spalax carmeli*, *Spalax galili*, and *Bos taurus*, Pregnancy-Specific Glycoproteins in *Homo sapiens* and *Mus musculus*, Pregnancy-Associated Glycoproteins in *Bos taurus*, genes related to the thymus-specific serine protease *PRSS16* (protease, serine 16) in *Loxodonta africana*, and the expansion of the growth hormone and chorionic gonadotropin beta gene families in anthropoid primates [8, 9]. Expression patterns of specific genes also can vary widely between species. For example, the expression of orthologous genes in the placenta have been found to vary substantially between *Homo sapiens* and *Mus musculus* [5].

**Figure 1:**
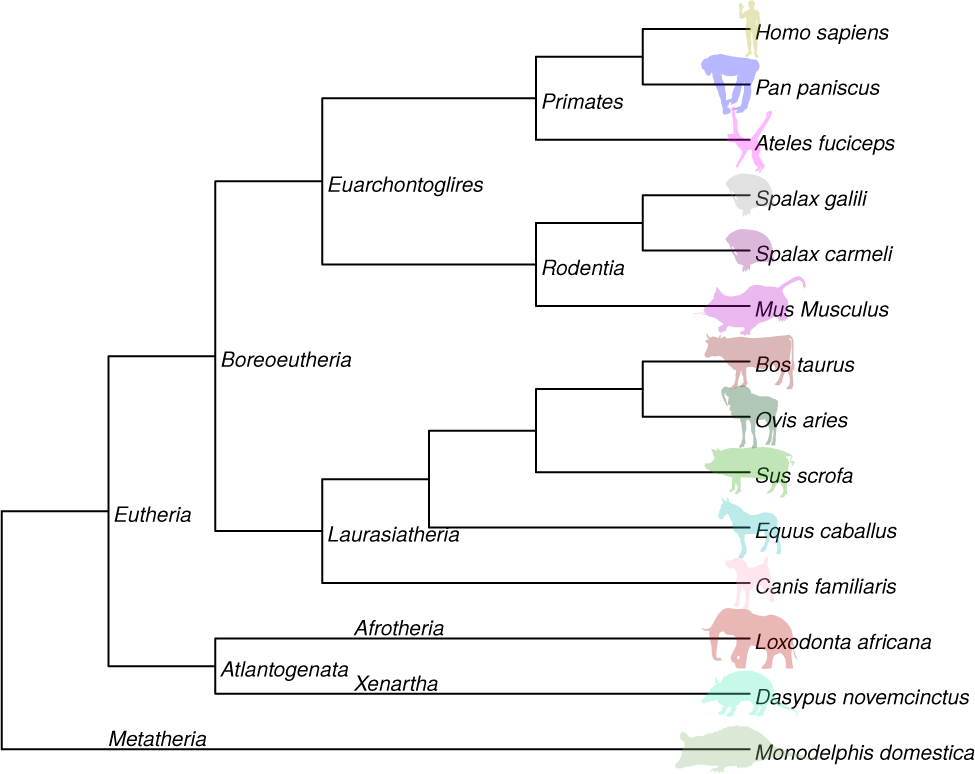
Species tree for organisms studied showing the relationship between species. Branch lengths are not to scale. Silhouettes of species were obtained from http://phylopic.org.

To address our questions we sequenced and quantified the total RNA of mammalian placentas at term, including *Homo sapiens* [human], *Pan paniscus* [bonobo], (the sister species to *Pan troglodytes*), *Ateles fusciceps* [spider monkey], *Mus musculus* [mouse], *Spalax galili* [blind mole rat from Upper Galilee], *Spalax carmeli* [blind mole rat from northern Israel], *Canis familiaris* [domestic dog], *Bos taurus* [domestic cow], *Loxodonta africana* [African savanna elephant], *Dasypus novemcinctus* [nine-banded armadillo], and *Monodelphis domestica* [gray short-tailed opossum]. These species represent four major branches of eutherian mammals (Euarchontoglires, Laurasiatheria, Afrotheria, and Xenartha) as well as marsupials (Fig. 1). These placentas also represent a wide array of placental morphologies as well as traits such as number and size of offspring and length of gestation (Supplemental Table S2). We use these data, along with existing RNA-Seq data from *Mus musculus* [10], *Homo sapiens* [11], *Equus caballus* [horse] [12], *Ovis aries* [13], and *Sus scrofa* [domestic pig] [14] to: 1) identify the non-housekeeping genes which are expressed in the placentas of all eutherian species analyzed and therefore represent the **core functionality of the placenta**; 2) identify genes whose expression level changes significantly in each of the placenta morphology types.

## 2. Methods

### 2.1. Collection of Placental Tissue and RNA Extraction

Fetal tissue was collected from the placenta of nine mammalian species (*Homo sapiens* [human], *Pan paniscus* [bonobo], *Ateles fusciceps* [spider monkey], *Canis familiaris* [domestic dog], *Bos taurus* [domestic cow], *Loxodonta africana* [African savanna elephant], *Monodelphis domestica* [gray short-tailed opossum], *Spalax galili* [blind mole rat from Upper Galilee], *Spalax carmeli* [blind mole rat from northern Israel] [15]; n=1 for all collected species). Information on specimen ID, number of specimens, and collector is shown for each sample in Supplemental Table S3. In the case of *Pan paniscus*, *Bos taurus*, and *Loxodonta africana*, the placenta was collected at term upon expulsion from the mother following delivery. Placental tissue from *Homo sapiens* was collected upon delivery via a cesarean section. Near-term placental tissue from *Canis familiaris* was collected upon the death of the mother and fetus from a car accident. Placental tissue from *Monodelphis domestica* was sampled at fetal stage 32, roughly one day prior to birth. Only the fetal portion of the placenta was sampled at the maternal-fetal interface (i.e. villous portion for *Homo sapiens*, *Pan paniscus*, *Canis familiaris*, and *Loxodonta africana*; cotyledon for *Bos taurus*; diffuse placenta for *Monodelphis domestica*). The placenta membranes were separated from the fetal portion by dissection, and the resulting fetal portion was washed twice in PBS. All tissues were collected and stored in RNAlater (Life Technologies, Carlsbad, CA) until RNA isolation could be performed. Tissues were homogenized in Trizol, and total RNA was isolated following the manufacturer’s instructions. RNA was further purified using RNeasy Midi Kit (Qiagen, Valencia, CA), in conjunction with the RNase-Free DNase Set (Qiagen, Valencia, CA), following the manufacturer’s protocol to remove any contaminating DNA from the sample. Total RNA was then evaluated on a Nanodrop 1000 (Thermo Scientific, Wilmington, DE) and an Agilent Bioanalyzer 2100 (Santa Clara, CA) to confirm concentration and quality of the isolation. RNA quality was assessed using A260/A280 and A260/A230 absorbance values.

### 2.2. Sequencing

All RNA-Seq mRNA isolation, library construction, and Illumina sequencing was performed by the Applied Genomics Technology Center (AGTC) at Wayne State University School of Medicine using 2.5-5 µg of total RNA and the standard Illumina protocol. Paired-end sequencing was performed using the Illumina Genome Analyzer II with an insert size of 230 base pairs (bp). Read lengths for species are shown in Supplemental Table S4.

#### 2.2.1. Additional Placenta Transcriptomes

For more complete evolutionary comparisons additional transcriptomes of placenta in *Mus musculus* [mouse] (n=17) [10], *Homo sapiens* (n=26) [11] *Equus caballus* [horse] [12], *Ovis aries* [sheep] [13], and *Sus scrofa* [pig] [14] were obtained from the Sequence Read Archive (Supplemental Table S3) and analyzed using the same methodology as RNAs sequenced specifically for this study.

### 2.3. Transcriptome Assembly, Alignment, and Quantification

Reads quality was evaluated using FASTQC v0.11.2 (http://www.bioinformatics.babraham.ac.uk/projects/fastqc/). Transcriptomes without a suitable reference genome (*Ateles fusciceps*, *Spalax galili*, *Spalax carmeli*; italic reference in Supplemental Table S4) were assembled using Trinity v2014-07-17 [16] and annotated using Diamond [17] against either the *Homo sapiens* (*Ateles fusciceps*) or *Mus musculus* (*Spalax galili* and *Spalax carmeli*) transcriptome (Supplemental Table S4), and the the match with the highest bitscore was chosen as the annotation. Transcripts where the highest scoring match had a bitscore below 40 were left un-annotated. All transcriptomes were aligned to the reference (or assembled) genome using STAR v2.4.2a [18] (Supplemental Figs. S1 and S2 and Table S4) and expression values in Fragments Per Kilobase of exon per Million reads (FPKM) were quantified using Cufflinks v2.2.1 [19] (Supplemental Tables S6 to S19). The genome of the closely-related *P. troglodytes* (separation ≈ 1 million years [20]) was used as a reference for *Pan paniscus*.

### 2.4. Comparative Analysis of Gene Expression

Orthologous pairs between *Homo sapiens* and all other species were identified using the Orthologous MAtrix project (OMA) database [21]. In cases where a gene did not have a human ortholog pair identified by OMA, Ensembl v.80 one2one orthologs obtained from Biomart using the biomaRt R package [22] were used to select the appropriate orthologous pair. A total of 5390 ortholog pair sets in all 14 species were expressed (FPKM *≥* 10) in at least one species. We also obtained orthology from the perspective of each species to all other species using the biomaRt package and the Ensembl v.80 database to determine the percentage of highly expressed genes that had an orthology of one-to-one, one-tomany, many-to-many, or novel/unknown. For future analyses by other investigators, we have also provided the human gene which most closely aligns to each expressed annotated protein-generating sequence of all species analyzed using Diamond [17].

Significant shifts in expression levels between groups of species were calculated using Student’s t test on ortholog pairs using the mean of each species to avoid large biases towards samples with *n ≫* 1 and considered significant if the False Discovery Rate (FDR) as estimated by the Benjamini and Hochberg procedure [23] was *≤* 0.05. For *Homo sapiens* and *Mus musculus* lineage specific calculations, a type-III anova was calculated using lme4 [24] with each species having separate random effects. Shifts in expression levels are reported in log_2_ Fold-Change (FC).

Significant shifts in expression levels which correlate with placenta morphology (shape, interdigitation, and intimacy) were identified using ANOVA on generalized linear models with the log FPKM as the response variable and the morphology as the predictor variable. Placenta morphology with a definable order (interdigitation and intimacy) were additionally analyzed using proportional odds logistic regression using the MASS package in R [25].

### 2.5. Gene Set and Pathway Enrichment Analysis

Gene sets and pathways in were obtained from MSigdb v5.1 [26] and Gene Ontology to form a superset of available pathway databases. We calculated the over-representation of gene lists in these gene sets using Fisher’s exact test and corrected for multiple testing by estimating the FDR using the Benjamini and Hochberg procedure [23].

To identify genes which have evidence of association with adverse pregnancy outcomes and other placenta-relevant terms of interest, we used the “pubmed_to_grid” component of Function2Gene [27] which conducts a term by gene search of PubMed results.

### 2.6. Sequences, FPKMs, and Code

All of the code used to produce every analysis presented in this paper and the manuscript itself is available at https://github.com/uiuc-cgm/placenta-viviparous.git. All of the sequences and assembled transcriptomes used are deposited in SRA, and new sequences for this work can be obtained under the series GSE79121 (Supplemental Table S3). The mean FPKM for gene in each species with ortholog annotations are located in the supplemental data file all_species_mean_fpkm.txt.xz and the per-sample FPKM values are in all_species_per_sample_fpkm.txt.xz.

## 3. Results

### 3.1. Transcriptome Statistics

Supplemental Fig. S1 shows the RNA-Seq read and alignment statistics for all species. For transcriptomes sequenced with 76 bp (*Bos taurus*, *Homo sapiens*, *Loxodonta africana*, and *Monodelphis domestica*),the number of trimmed reads ranged from 1.1×10^6^ to 3.7×10^7^ with an average of 88.8% of reads mapping to the respective reference genome (Table S4). For transcriptomes sequenced with 150 bp (*Canis familiaris*, *Ovis aries*, and *Pan paniscus*), the number of trimmed reads ranged from 3.7×10^7^ to 8.4×10^7^ with an average of 92.8% of reads mapping to the respective reference genome (Supplemental Table S4). For transcriptomes sequenced with 100 bp (*Ateles fusciceps*, *Dasypus novemcinctus*, *Equus caballus*, *Spalax carmeli*, and *Spalax galili*), the number of trimmed reads ranged from 1.9×10^7^ to 1.9×10^8^ with an average of 92.4% of reads mapping to the respective reference genome (Supplemental Table S4). For transcriptomes sequenced with 35 bp (*Mus musculus*), the number of trimmed reads ranged from 2.8×10^6^ to 5.2×10^6^ with an average of 89.6% of reads mapping to the respective reference genome (Supplemental Table S4). Read length was not a significant predictor of unmapped read percentage (Supplemental Fig. S2). Between 20% and 37% of all highly expressed genes in each species were ortholog pairs with all other analyzed species, with the remainder being mostly one-to-many or many-to-many in comparison to at least one other species (Supplemental Table S5). The expression of all genes with FPKM *≥* 50 is shown in Supplemental Tables S6 to S19.

We performed gene set enrichment analysis on both the top 1% of transcripts by FPKM (Supplemental Tables S20 to S33) and the top 100 transcripts by FPKM (Supplemental Tables S34 to S47); common processes include protein transport and viral transcription/expression (Supplemental Table S48).

### 3.2. Homo sapiens Lineage Specific Genes and Their Association with Pre-eclampsia

Using a type-III ANOVA to account for repeated testing of *Homo sapiens* (see Section 2.4 for details), we identified multiple genes which are differentially expressed in the *Homo sapiens* lineage in comparison to all other lineages (Table 1, Supplemental Table S49). These genes include multiple genes which are thought to be involved in pre-eclampsia. Notably, *CRH* (corticotropin releasing hormone) (FDR=2.1×10^−5^, FC=9.6, Supplemental Fig. S3), has been shown to be more highly expressed in placentas from pre-eclamptic pregnancies [28, 29], where it is hypothesized to activate decidual macrophages. Likewise, *KISS1* (KiSS-1 metastasis-suppressor) (3.1×10^−5^, FC=9.3, Supplemental Fig. S4) also shows increased expression in pre-eclamptic placentas, and is involved in modulating trophoblast invasion [30]. Two other genes, *PAPPA* (pappalysin 1) (3.7×10^−9^, FC=8.5 Supplemental Fig. S5) and *ADAM12* (ADAM metallopeptidase domain 12) (1.9*×*10^−10^, FC=8.5, Supplemental Fig. S6) are also differentially expressed but are negatively correlated with pre-eclampsia [31].

**Table 1.**
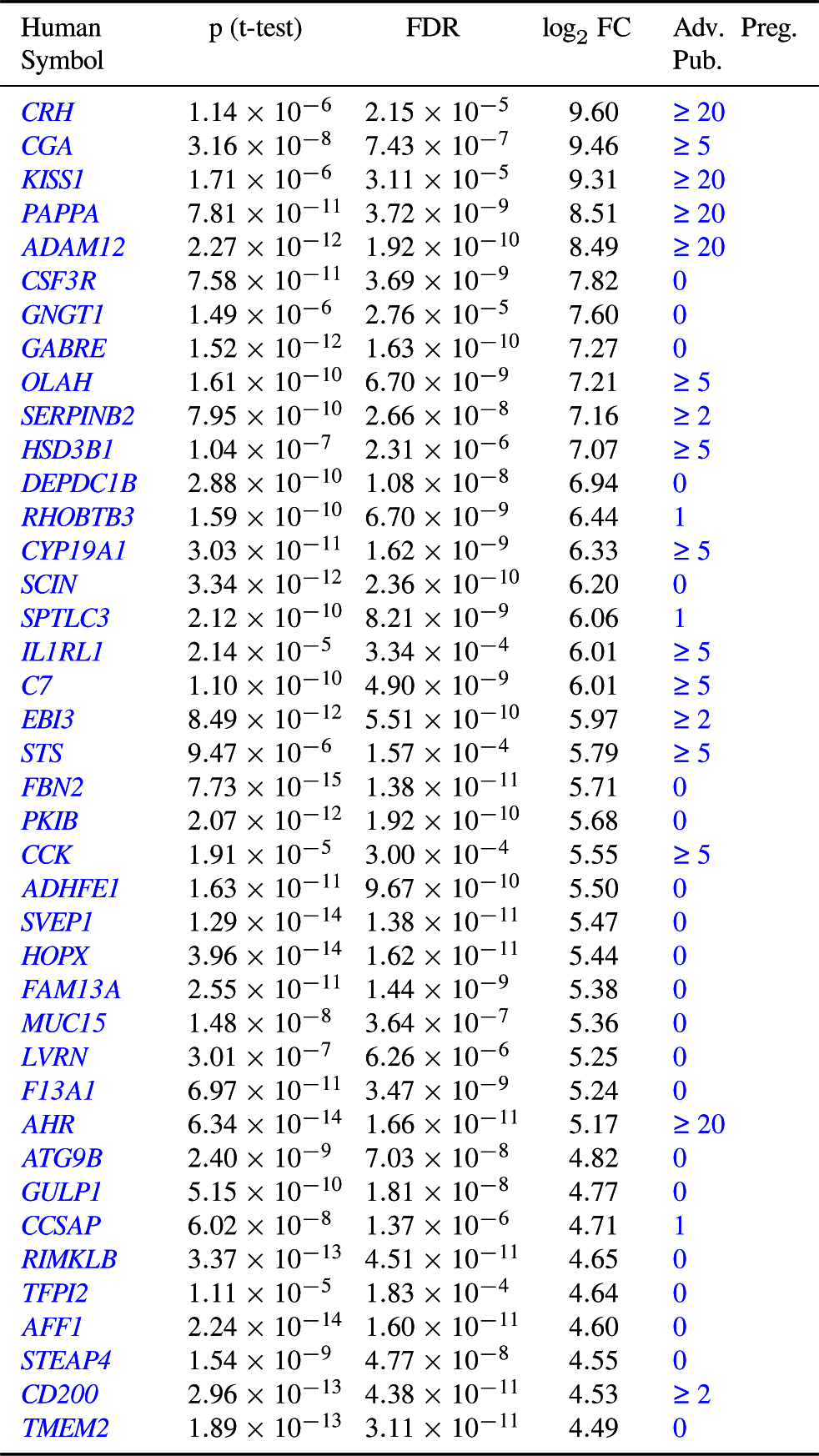
Genes whose expression changed in *Homo sapiens* as compared to all other analyzed species. A nested type-III anova was used to calculate p values, and the top 40 genes by log_2_ Fold-Change which changed significantly (FDR *≤* 0.05) are shown. FDR is the False Discovery Rate as estimated by the Benjamini and Hochberg procedure [23]. log_2_ FC is the base-2 logarithm of the fold change from *Homo sapiens* to all other analyzed species. The human symbol links to a brief explanation of the gene function in Ensembl. Adv. Preg. Pub. is the approximate number of publications indexed by PubMed mentioning this gene and an adverse pregnancy outcome (pre-eclampsia, premature labor, gestational diabetes, intrauterine growth restriction, or premature birth) as of February 2017, and links to the most current PubMed search results.

### 3.3. Other Lineage and Clade Specific Genes

We also compared differences in the *Mus musculus* lineage in comparison to all other lineages using a type-III ANOVA as in the *Homo sapiens* analysis, and found few genes which are highly differentially expressed and are also associated with adverse pregnancy outcomes, including *APOB* (apolipoprotein B) and *AFP* (alpha fetoprotein) (Supplemental Table S50). Comparisons between rodentia (*Mus musculus*, *Spalax galili*, and *Spalax carmeli*) and all other species using the mean of each species expression did not indicate any significant differences (Supplemental Table S51) nor did the equivalent comparisons between laurasiatheria and all other species (Supplemental Table S52) and laurasiatheria and euarchontoglires (Supplemental Table S53). Additional placenta samples of each species and the collection of the placentas of additional laurasiatherians may resolve the lack of significance due to insufficient power.

### 3.4. Core Placenta Transcriptome

The 115 genes comprising the core placenta transcriptome are shown in Fig. 2 ordered by their median transcription level in FPKM. We considered a gene to be core to the placenta transcriptome if an ortholog pair of that gene was expressed with FPKM *≥* 10 in all eutherian species studied (excluding the metatherian, *Monodelphis domestica*) and was not a human housekeeping gene according to Eisenberg & Levanon [32]. Genes which meet the FPKM criterion and are housekeeping genes are depicted in Supplemental Fig. S7. Figure 3 shows the number of genes which would be included in the core placenta set if the FPKM cutoff was set differently; for example, choosing an FPKM cutoff of 1 yields 1571 genes in the core transcriptome. In contrast to the shape of the number of genes included in the core placenta transcriptome, the set of *Homo sapiens* housekeeping genes expressed all placentas is flatter (Supplemental Fig. S9), and includes more of the total fraction of genes (0.6 vs 0.4). An alternative approach to identify housekeeping genes is to use the tissue specificity index [33]. Supplemental Table S54 lists the tissue specificity index for all housekeeping and non-housekeeping genes; housekeeping genes expressed in all placentas have a lesser median tissue specificity index than non-housekeeping genes (Supplemental Fig. S8). We also note that housekeeping genes have been defined in *Homo sapiens*, and may not be housekeeping genes in other species; generating tissue expression atlases and examining the evolution of expression in non-model organisms is a promising area of future research.

**Figure 2:**
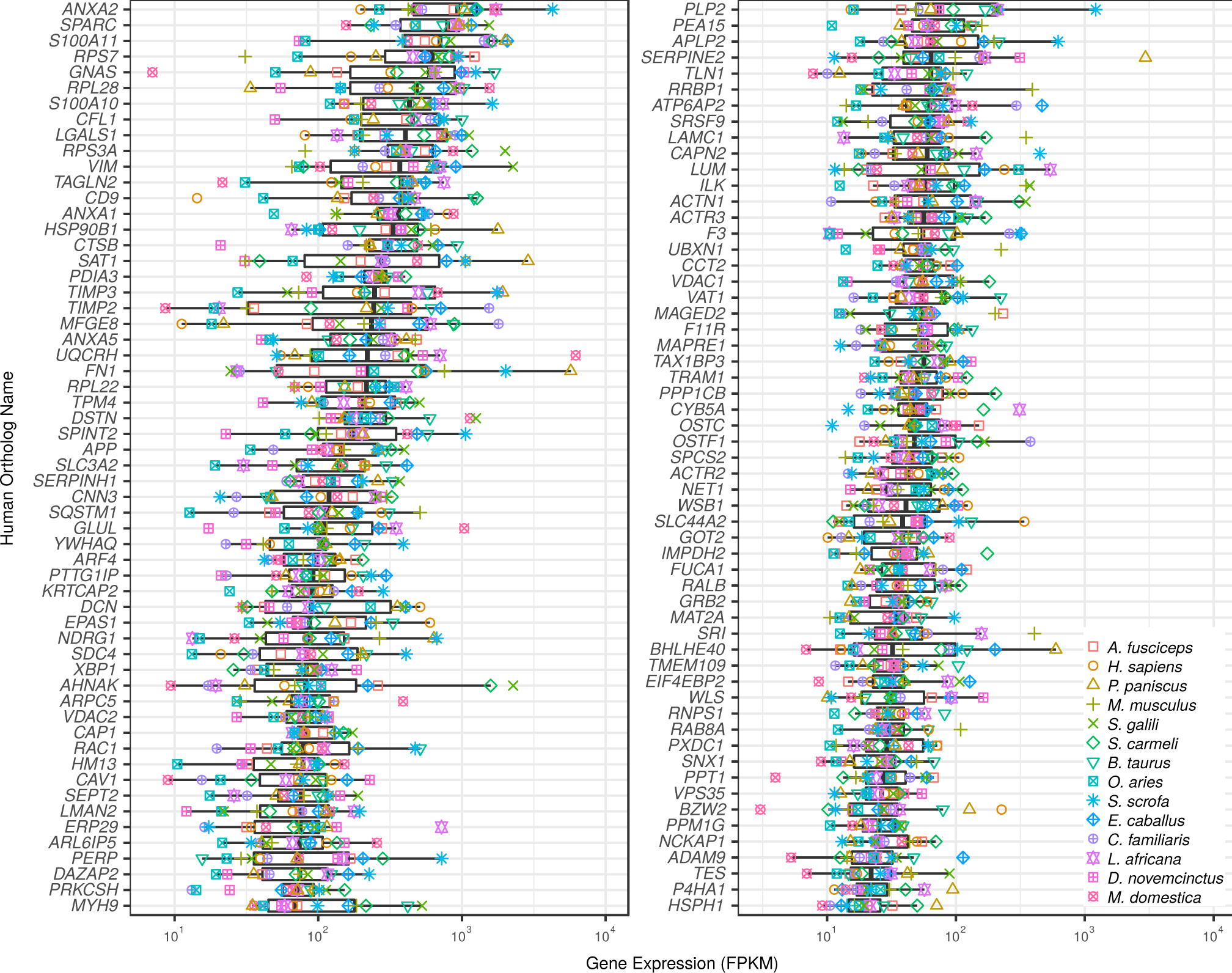
Core Placenta Transcriptome. 115 ortholog pairs of *Homo sapiens* genes expressed (FPKM*≥* 10) in the placenta of all 13 eutherians studied with housekeeping genes [32] removed ordered (from top to bottom, left to right) by median expression. If we also require expression (FPKM*≥* 10) in the marsupial, *Monodelphis domestica*, there are 95 genes which meet this criteria. Figure 3 shows the shape of the distribution of the number of genes in the core placenta transcriptome if different FPKM and species restriction criteria are utilized.

**Figure 3:**
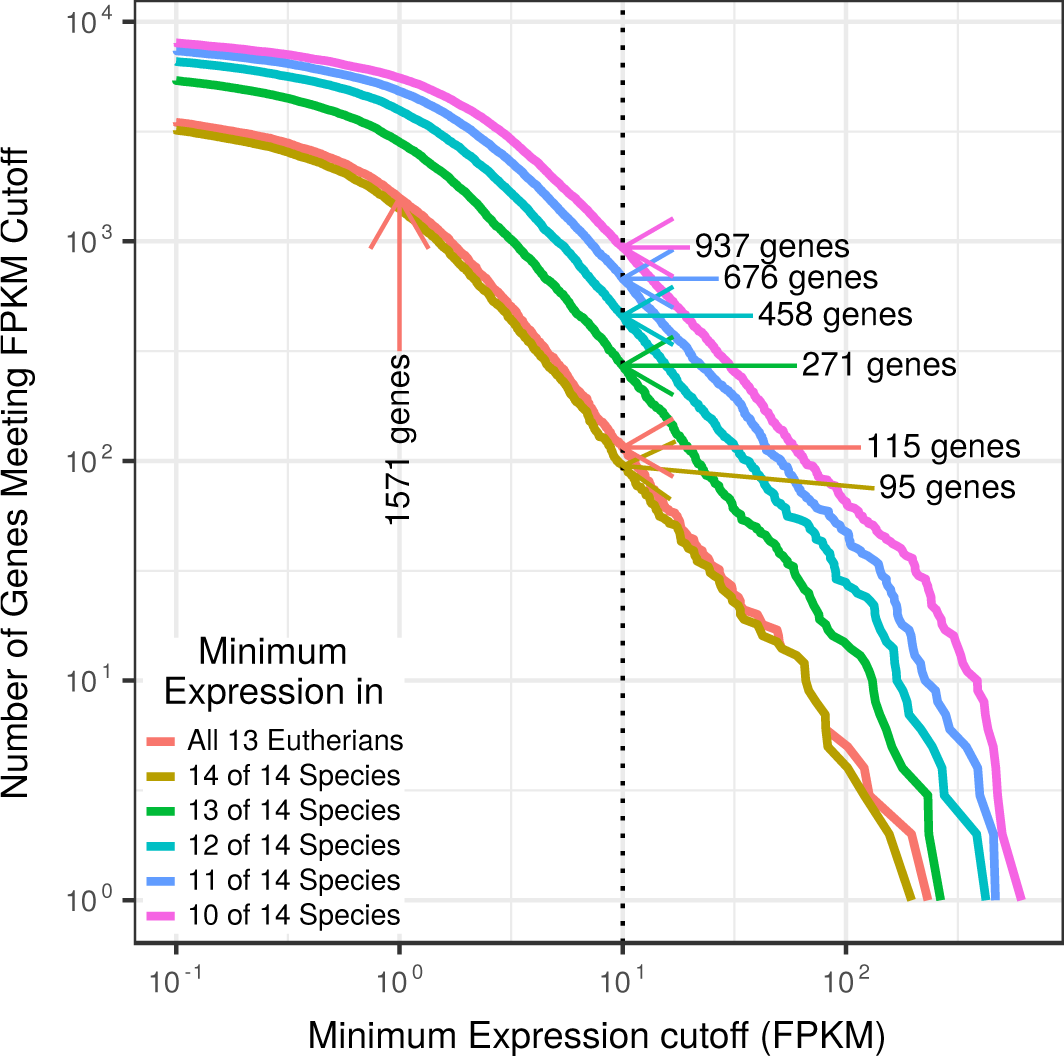
Distribution shape of placenta core genes for all possible number of genes. The vertical axis shows the number of ortholog pairs included in the core placenta transcriptome (for all possible number of genes) if all 13 eutherians (or 14 of 14 species, or 13 of 14 species, etc.) have a minimum expression in FPKM equal to the horizontal axis. For example, if the cutoff is set to 10 (vertical black line), 115 genes would be included if the expression of a set of ortholog pairs of all 13 Eutherians must be at 9 least 10 FPKM, 271 ortholog pairs would be included if 13 of 14 species have an expression of at least 10 FPKM, and 937 genes would be included if 10 of 14 species must have an FPKM of at least 10.

Supplemental Table S55 lists the significant enrichment of Gene Ontology terms in the core placenta transcriptome; notable terms include those involved in cytoskeletal structure, protein transport, membrane protein structure, and inflammation resolution. Increasing the permissiveness of the core placenta transcriptome by only requiring 10 of 14 species to express an ortholog pair with FPKM *>* 10 and results in the same terms also being significantly enriched, along with additional terms (Supplemental Table S55). Supplemental Table S56 shows the pathways with significant and suggestive enrichment in the core placenta transcriptome.

If we use gene trees to define the core placenta transcriptome instead of ortholog pairs (where the mean expression for genes in the tree which are present must be FPKM *≥* 10), we find 71 unique gene trees which include most (68%) of the genes present in the core placenta transcriptome (Supplemental Figs. S10 and S11). Relaxing the mean gene tree expression requirement to FPKM *≥* 5 includes nearly all of the genes (91.3%).

#### 3.4.1. Annexin Complexes

Multiple components of annexin complexes are included in the core transcriptome, including *ANXA1* (annexin A1), *ANXA2* (annexin A2), *S100A10* (S100 calcium binding protein A10 (calpectin-I light polypeptide)), and *S100A11* (S100 calcium binding protein A11 (calgizzarin)). ANXA1 is involved in the resolution of the inflammation response through inhibition of phospholipase A2 activity and signaling through the formyl peptide receptor family, and has previously been hypothesized to be important for the maintenance of the anti-inflammatory state during pregnancy [34]. ANXA2 is known to interact with S100A4, S100A6, S100A10, and S100A11 [35], and there is evidence for its interaction with S100P as well; it is involved in cell-cell interaction as well as vesicle trafficking and von Willibrand Factor secretion. The expression regulation of another core gene, *TIMP3* (TIMP metallopeptidase inhibitor 3), has been implicated in pre-eclampsia [36].

Another annexin which is highly expressed in all term placentas is *ANXA5* (annexin A5), which was originally identified as an anti-coagulant protein, but is also involved in syncytium formation, membrane repair, and reduction of spontaneous abortion [37]. Other notable syncytiotrophoblast fusion proteins, Syncytin-1 (*ERVW-1* (endogenous retrovirus group W member 1)) and −2 (*ERVFRD-1* (endogenous retrovirus group FRD member 1)) were not present in the core placenta, as they (or genes similar to them) are not highly expressed in *Canis familiaris*, *Pan paniscus*, or *Ovis aries*, nor was the Syncytin-2 receptor, *MFSD2A* (major facilitator superfamily domain containing 2A) (FPKM *≥* 10 only in *Homo sapiens*, *Mus musculus*, and *Canis familiaris*).

#### 3.4.2. Invasion: EGF and Actin Y in the Core Placenta

The core placenta transcriptome is significantly enriched in genes which are involved in multiple biological pathways (Supplemental Table S56). The most abundant set of significantly enriched molecular pathways are related to the migration of the trophoblast into the endometrium, including the EGFR1 pathway (FDR = 6.5×10^−4^), Focal Adhesion pathway (FDR = 6.5×10^−4^) and the Actin Y pathway (FDR = 9.2×10^−3^). Two other significant pathways, the mPR/Progesterone pathway and Rho cell motility pathways, feed into the Actin Y pathway as well. Supplemental Fig. S12 shows the EGF pathway in humans with the median expression in FPKM of genes in the pathway shown. Notably, many of the genes on the GRB2→MYC pathway are strongly expressed; their activation leads to an increase in migration and invasion in response to EGF binding to EGFR1 [38]. The Actin cytoskeleton pathway is depicted in Supplemental Fig. S13, which shows that *ITGB1* (integrin subunit β 1) and its binding protein *ITGB1BP1* (integrin subunit β 1 binding protein 1) as well as *FN1* (fibronectin 1) are highly expressed, which activates Rac. This potentially leads to the formation of focal adhesions and lamellipodia formation through inhibition of GSN and HSPC300. Other components of cell-cell adhesion, including *F11R* (F11 receptor) which forms tight junctions in epithelia, the proteoglycans *SDC4* (syndecan 4) and *LUM* (lumican), and *LAMC1* (laminin subunit γ 1).

### 3.5. Placenta Morphology: Shape

Supplemental Table S57 shows the 50 genes which are most significantly associated with the shape of the placenta, as well as the shape in which the difference is most significant. The most significant genes which were associated with diffuse placenta shape include *EREG* (epiregulin), a ligand of EGFR, *UNC93A* (unc-93 homolog A (*C. elegans*)), a gene similar to *UNC93B1* (unc-93 homolog B1 (*C. elegans*)) which is involved in the balancing between Tlr7 and Tlr9 immune response in mice [39], and *WNT5A* (Wnt family member 5A), which is a member of the Wnt ligand family whose signaling pathway has been implicated in motility in human trophoblast cells [38].

### 3.6. Placenta Morphology: Degree of Intimacy and Interdigitation

For the degree of intimacy and interdigitation, we saw multiple genes which were associated with different levels of interdigitation and intimacy (Supplemental Tables S58 and S60) using ANOVA, but when we used an ordered factor analysis (proportional odds logistic regression), none of these genes were retained (Supplemental Tables S59 and S61). We suspect that this is due to the fact that the ancestral state of all eutherians is a labyrinthine, hemochorial placenta [40]. This implies that the transcriptome of intermediately invasive or interdigitated placentas need not be intermediate between the transcriptomes of less intimate (folded/lamellar) or interdigitated placentas (epitheliochorial) and most intimate (hemochorial) or interdigitated placentas (labyrinthine) because these morphologies have evolved independently. An alternative is that the limited number of species with placentas of intermediate interdigitation and intimacy in this study limits the ability to detect genes which whose expression correlates with degree of intimacy and interdigitation. Future analyses of additional placentas with intermediate interdigitation and intimacy should better illuminate the pathways which are responsible for these placental morphologies.

## 4. Discussion

### 4.1. Homo sapiens Specific Expression Shifts and the Evolution of Pre-eclampsia

The *Homo sapiens* placenta is distinctive from many other primates in that it possesses a deeply invasive trophoblast that penetrates the inner myometrium of the uterus [41]. The near exclusivity of deep trophoblast invasion in humans has also led many to speculate on the origin of various disorders, which are almost exclusively symptomatic of human pregnancy. For example, pre-eclampsia is a human disease that occurs in 4% of pregnancies in the United States [42]. Pre-eclampsia can be attributed to the lack of adequate invasion of maternal tissue, and is noted by the onset of hypertension and proteinuria (or in the absence of proteinuria, thrombocytopenia, impaired liver function, renal insufficiency, pulmonary edema, or cerebral or visual disturbances) [42]. Some cases of pre-eclampsia can develop into eclampsia, a more severe condition resulting in seizures during pregnancy and death. Lack of invasion in pre-eclampsia may be attributed to the inefficiency of growth regulators as well as the inability to sufficiently suppress the maternal immune response [34]. Although there are speculative cases of eclampsia in great apes, it is exceedingly rare in these species [41].

The identification of multiple genes (*CRH*, *ADAM12*, *KISS1*, *PAPPA*, *IL1RL1* (interleukin 1 receptor like 1), and *PLAC1* (placenta specific 1); Table 1) which are associated with pre-eclampsia which have increased expression on the *Homo sapiens* lineage may explain the larger proportion of pre-eclampsia observed in humans. However, there are still genes which are associated with pre-eclampsia which are also more highly expressed in the *Mus musculus* lineage such as *Afp* (α-fetoprotein) and *Gjb2* (gap junction protein β 2). Furthermore, the biological processes which ultimately result in pre-eclampsia occur well before term; all of our placentas are term placentas and provide limited information about the transcriptome during the onset of pre-eclampsia.

### 4.2. Core placenta transcriptome

The core placenta transcriptome is enriched in multiple genes which are critical for the function of the placenta, including vesicular transport, immune regulation, cell fusion and membrane repair, and invasion (Supplemental Tables S55 and S56).

#### 4.2.1. Immune Regulation and Annexins

The transcriptomes from multiple different species enabled us to identify the set of expressed genes which are common to all placentas. Components of the annexin complexes, including *ANXA2*, *ANXA1*, *S100A11*, and *S100A10* are expressed in all placentas examined, and argue for the evolutionary importance of annexins in placenta function, where they likely establish maternal-fetal tolerance[34, 35, 43]. We suggest that further research into the molecular role of these annexin complexes may illuminate how immune tolerance is maintained during placentation, because annexins appear to be the major class of immune modulatory transcripts which are expressed in all placentas analyzed.

#### 4.2.2. Formation of the syncytiotrophoblast

The evolutionarily convergent capture of endogenous retroviruses to form expressed fusogenic proteins such as Syncytin-1 and −2 (*ERVW-1* and *ERVFRD-1*, respectively) has been implicated in syncytiotrophoblast formation in *Homo sapiens* and many other species [9]. While genes which are similar to *ERVFRD-1* and *ERVW-1* are expressed in most of the placentas analyzed here, there are no highly expressed genes with sequence similarity in *Pan paniscus* and *Canis familiaris*, even though these mammals form syncytiotrophoblasts [44]. Because the formation of syncytiotrophoblast is the stem state of Eutheria, Afrotheria, Euarchontoglires, and Laurasiatheria [44], and the syncytiotrophoblast must be continuously replenished, the gene(s) responsible for cytotrophoblast fusion to form the syncytiotrophoblast should be highly expressed in the core placenta. *ANXA5* is known to be expressed at multiple stages of placentation, and is capable of repairing membrane defects and promoting the fusion of cells [45] in the presence of Ca2+. We hypothesize that *ANXA5* is essential for the completion of fusion and the maintenance of membrane repair during the flux of nutrients and waste in the placenta [37, 45].

#### 4.2.3. Invasion: EGF and CD9 in the Core Placenta

The high expression of components of the EGFR signaling pathway (Enrichment FDR = 6.5×10^−4^) in the core placenta transcriptome (including *GRB2* (growth factor receptor bound protein 2), *RALB* (RAS like proto-oncogene B)) as well as annexins which are involved in the spatio-temporal regulation of EGF receptor signal transduction (*ANXA1*, *ANXA2*, and *S100A11* [46]) in the core placenta is consistent with previous findings showing that alterations to the EGF signaling system change invasion of trophoblast cells into the spiral arteries [47]. Another component of the core placenta transcriptome, *CD9* (CD9 molecule), has been shown to suppress *Homo sapiens* extravillous trophoblast invasion [48], and we hypothesize that it is responsible for maintaining the balance between invasion and attachment in the other placentas examined here as well.

### 4.3. Limitations

This study includes only term placentas, and our conclusions are limited to the set of genes which continue to be expressed at term. We suspect that many genes which are responsible for establishing shape, intimacy, and interdigitation are expressed and function at earlier time points where placental morphology is being established. Furthermore, while these samples have been dissected to only include the fetal side, they likely include substantial cell and tissue heterogeneity. We are therfore not yet able to assign specific genes in our core placental transcriptome to specific cell types and tissues. We believe that samples of placentas from earlier in gestation and from multiple locations within the placenta and at multiple will address these limitations.

### 4.4. Conclusion

We analyzed the sequenced transcriptome of 55 placentas from 14 species spanning the phylogenetic tree of mammals. This revealed both lineage-specific differential expression of gene families as well as a set of core, non-housekeeping genes which are critical for the proper organization and functioning of the placenta.

The core placenta transcriptome contains genes which are involved in maternal-fetal tolerance and genes responsible for cell-cell and cell-matrix interactions, such as the annexins and *LGALS1* (galectin 1). These genes were almost certainly present in the placenta of the most recent common mammalian ancestor of the species included in this study. We hypothesize that modifications to the expression or function of genes in the core placenta transcriptome will compromise the development and organization of the placenta.

We also identified genes differentially expressed on the *Homo sapiens* lineage which are implicated in pre-eclampsia, including *ADAM12*, *CRH*, and *KISS1*.

This study has identified many tantalizing molecular pathways which illuminate the vast diversity of placental morphology and may explain some placental malfunctions in *Homo sapiens*. However, the strength of our hypotheses are limited by our limited data; more placentas from multiple members of diverse clades and different time points are still needed to come to a more complete understanding of the genomic and molecular basis for placenta morphology and function. We look forward to collecting and analyzing such a dataset.

## Author’s contributions

Don Armstrong analyzed all of the data and wrote the paper.

Michael McGowan wrote the paper and performed initial analyses.

Amy Weckle performed all of the RNA extractions and managed the samples.

Priyadarshini Pantham wrote the paper and performed gene enrichment analyses.

Jason Caravas performed initial analyses.

Dalen Agnew provided samples.

Kurt Benirschke provided samples.

Sue Savage-Rumbaugh provided samples.

Eviatar Nevo provided samples.

Chong J. Kim provided samples.

Günter P. Wagner provided samples and wrote the paper.

Roberto Romero provided samples and conceived the study.

Derek E. Wildman conceived the study, wrote the paper, and performed analyses.

## Acknowledgements

The authors would like to specially acknowledge those groups whose publicly available data contributed to the completeness of this work, as well as the suggestions of the anonymous reviewers which improved this paper and the advice of Monica Uddin in preparing this manuscript.

## Funding

Partial support for this research came from the *Eunice Kennedy Shriver* National Institute of Child Health and Human Development (NICHD), National Institutes of Health, U.S. Department of Health and Human Services (N01-HD-3342), the National Science Foundation (BCS-0827546 and BCS-0751508), and a seed grant from the Institute for Genomic Biology at the University of Illinois, Urbana-Champaign.

